# Unsaturated fatty acids profiling in live *C. elegans* using real-time NMR spectroscopy

**DOI:** 10.1101/2021.04.02.438181

**Authors:** Bruno Hernández Cravero, Gastón Prez, Verónica A. Lombardo, Andrés Binolfi, Diego de Mendoza

## Abstract

Unsaturated fatty acids (UFAs) impact central cellular process in animals such as membrane function, development and disease. Perturbations of UFAs homeostasis contribute to the onset of metabolic, cardiovascular and neurodegenerative disorders. Nevertheless, links between lipid desaturation fluctuations and the dynamics of mono and polyunsaturated fatty acid synthesis in live animal physiology are poorly understood. To advance in the understanding of this process, we decided to study *de novo* UFAs synthesis with the highest resolution possible in live *Caenorhabditis elegans*. Conventional lipid analysis in this organism involves solvent extraction procedures coupled with analytical techniques such as chromatography and/or mass spectrometry. These methodologies are destructive and prevent the access of information, linking *in vivo* UFA dynamics and functionality. To overcome these limitations, we used uniform ^13^C isotope labeling and real-time 2D heteronuclear NMR spectroscopy in live *C. elegans* to identify their UFA compositions and the dynamic response of these fatty acids during cold adaptation. Our methodology allowed us to monitor in real time the upregulation of UFA synthesis when ambient temperature is decreased. The analysis of UFAs synthesis in worms lacking the adiponectin receptor AdipoR2 homolog PAQR-2 during a temperature drop supports the pivotal role of this protein in low temperature adaptation and survival. Our results provide new insights about the environmental regulation of UFAs and establish methodological benchmarks for future investigations of fatty acid regulation under experimental conditions that recapitulate human diseases.

## INTRODUCTION

Unsaturated fatty acids (UFAs) are important membrane components and precursors of signaling molecules. The specific composition of subcellular membranes is of paramount importance for many cellular processes ranging from vesicular trafficking to organelle homeostasis and receptor signaling^1^. For example, recent studies suggest important roles for long chain monounsaturated and polyunsaturated fatty acids (MUFAs and PUFAs) in membrane deformation^2^, domain stability^3^, membrane fission events^4^ and membrane fluidity^5^. Therefore, many disease states are associated with membrane composition defects. UFAs have been directly linked to the onset of severe metabolic^6^, cardiovascular^7^ and neurodegenerative disorders^8^. Given its central and far-reaching importance, it is surprising that so little is known about the molecular mechanisms that regulate membrane composition and fluidity in animals.

The nematode *Caenorhabditis elegans* is a powerful model organism to study functions of MUFAs and PUFAs. The development of highly sensitive chromatographic and spectroscopic methods has enabled comprehensive lipid analysis in this system^9^. The simple anatomy and wide array of forward and reverse genetic engineering tools available in *C. elegans* makes this model ideal for discovering new roles and regulation for lipid metabolism^10^.

*C. elegans* are capable of synthesizing all necessary fatty acids *de novo*, and the core enzymes of fatty acid biosynthesis are conserved, including a range of fatty acid desaturase and elongase activities, enabling *C. elegans* to synthesize 20 carbon omega 6 and omega 3 PUFAs, as arachidonic (AA, 20:4n6) and eicosapentanoic (EPA, 20:5, n3) acids, respectively (**Supplementary Figure 1**)^10^. Studies in *C. elegans* have characterized numerous pathways that modulate lipid metabolism and fat storage including insulin/IGF, serotonin and nuclear receptor signals^10^. Furthermore, recent studies uncovered the adiponectin homolog PAQR-2 that together with the nuclear receptor NHR-49 and the Δ9 stearic CoA desaturase FAT-7 form a cold adaptation pathway in larva at low temperature^11–13^. Despite PAQR-2 being essential for low temperature adaptation in larva^12,13^, the pathway(s) promoting the expression of fatty acid desaturases by this sensor are not completely characterized.

To this day, conventional lipid analysis is commonly performed by solvent extraction-based methods coupled to gas chromatography/mass spectrometry (GC/MS)^9^, which are typically destructive and time consuming. Moreover, information linking cellular dynamics and functionality is often lost during extraction steps making them less useful for understanding the fundamental biological processes that are needed for controlling UFA homeostasis. Although fixative stains such as Oil Red O measure distributions of lipid-like species across tissues^14^, these fluorescent probes cannot provide detailed chemical information about fatty acid compositions, degrees of unsaturation and chain lengths. Recently, coherent Raman spectroscopy was used to study the distributions of lipid-rich deposits in live worms but information about individual lipid compositions was not accessible^15^. Thus, studying UFA compositions at high resolution in living cells and multicellular organisms is difficult to accomplish with existing methods.

Unlike other analytical tools, solution state NMR spectroscopy is non-destructive and enables high resolution monitoring of dynamic changes of biomolecules, including metabolites^16^, proteins^17^ and nucleic acids^18^ in a tag-free manner and in complex environments that range from cell extracts to whole cells. In particular, NMR metabolomics has emerged as a powerful tool to characterize the metabolic compositions of such complex samples, including animal tissue extracts or body fluids and their response to environmental or genetic conditions that result in metabolic changes ^19–23^. These strategies rely on the exquisite sensitivity of the NMR chemical shifts that enable the identification and quantification of individual metabolites in biological samples. Efforts in this field resulted in the construction of comprehensive databases that contain annotated chemical shifts for many biological molecules, thus greatly aiding their identification in newly analyzed samples^24^. Notably, these methods have been successfully applied to perform high-resolution metabolic studies *in vivo*. By introducing ^13^C-isotope enrichment in live multicellular organisms, such as the water flea *Daphnia magna*, metabolic compositions were analyzed using 2D ^1^H-^13^C correlation NMR spectra, thus expanding the number of compounds that can be identified and analyzed compared with standard 1D ^1^H NMR metabolomics studies^25,26^. Moreover, a continuous NMR flow system was used on ^13^C-enriched water fleas to assess the effects of growth conditions or stressors on organismal metabolism in real time^27,28^. Other applications of high resolution NMR studies in live animals include the monitoring of enzymatic reductions of exogenously delivered, ^15^N isotopically enriched, oxidized methionine derivatives in zebrafish embryos (*Danio rerio*), a widely used vertebrate model system for biological studies^29^. Finally, Park and co-workers recently applied NMR spectroscopy on live, ^13^C-isotopically enriched *C. elegans*, to characterize time-dependent metabolic changes of N2 and mutant worms with impaired AMP-activated kinase (AAK) activity, a central regulator of cellular metabolism^30^. Altogether, these studies demonstrate the advantages of NMR spectroscopy to perform high resolution metabolic NMR studies in live multicellular organisms.

Here, we employ uniform ^13^C isotope labeling followed by multidimensional real-time NMR in live *C. elegans* to identify UFA compositions and dynamic responses to temperature changes, an environmental cue that modifies relative UFA amounts by activating lipid desaturating enzymes. We compared N2 and *paqr-2* mutant worms and determined how PAQR-2-signaling influences this central aspect of UFAs homeostasis in live worms. Overall, we provide a methodological framework for studying the dynamics of fatty acids desaturation in live *C. elegans*, which opens new research avenues to characterize genetic, environmental, dietary and pharmacological effects on UFA composition and biosynthesis.

## MATERIALS AND METHODS

### Reagents

All buffers, salts and chemical were reagent grade and were used without further treatments. Unless specified, all reagents were purchased from Merck or Sigma.

### Nematode strains

*C. elegans* strains used in this work were obtained from the *Caenorhabditis* Genetics Center, University of Minnesota (https://cgc.umn.edu/): Bristol N2 (wild-type) and the single mutants *fat-3(ok1126),fat-4 (ok958), paqr-2 (tm3410)*.

### C.elegans maintenance and handling

Worms were routinely propagated on nematode growth medium (NGM) agar plates seeded with *E. coli* OP50 according to standard procedures^31^.

### Sample preparation for NMR experiments with live worms

To obtain uniformly ^13^C-isotopically enriched *C. elegans* samples, agar plates were spread using 10X pellet of an overnight culture of *E. coli* NA22^32^ grown in a M9 minimal medium supplemented with 2g/L of ^13^C-D-glucose (Cortecnec, France) and 1 g/L of ^14^N-ammonium chloride (Merck) as sole carbon and nitrogen sources, respectively. Typically, each worm NMR sample requires 50 mL of a saturated, ^13^C-isotopially-enriched, NA22 culture. Next, 60000 L1s worms from eggs prepared by bleaching^33^ with sodium hypochlorite treatment were grown at 25°C or 15 °C on 15 mL NGM plates seeded with 1 mL concentrated *E. coli* NA22 to L4 state. Non-isotopically enriched NMR samples were prepared using *E.coli* OP50 cultured in LB-broth. Cold-sensitive *paqr-2* worms were grown similarly at 25°C. For *paqr-2* experiments performed at 15°C we added 0.05% of Nonidet P-40 to the culture media following previously established procedures^12^.

Synchronized L4 worms were collected in a 15 mL falcon tube, centrifuged for 2 min at 2000 g to remove media and washed twice on M9 buffer to remove bacteria. We then discarded the supernatant, added M9 buffer to complete a volume of 450 μL and supplemented this solution with 50 μL of D_2_O (99.9 %). Worms were gently resuspended with a cut pipette tip, loaded in a 5 mm NMR Shigemi tube and decanted to the bottom by gentle shaking. We perform the NMR experiments with 40000-60000 L4 worms without applying the tube plunge.

### C. elegans lifespan analysis

Worms lifespan during NMR acquisitions was assessed on paralleled, independent samples by removing small aliquots of the worm suspension from the NMR tube at 2, 4, 6 and 8 hs after the experiments started. Worms were then transferred to fresh NGM plates where we counted the live and dead ones to obtain the survival percentages at each time point. Viability was determined by spontaneous and/or touch-induced movement. All observations were performed on an Olympus MVX10 with 10X and 40X magnifications. Images were taken with an Olympus DP72 camera. Each assessment was done in triplicates.

### Lipid Extraction

To extract lipids from whole worms, we used an adaptation of the protocol reported by Folch and co-workers^34^. Briefly, lipid extracts were made from approximately 200 mg of frozen N2, *fat-3* or *fat-4* worm pellets grown at 20 °C. Pellets were initially washed with M9 buffer, resuspended in 1.3 ml pure methanol and sonicated on ice one time for 30 seconds at maximum power (Branson Sonifier). We waited for 2 min between each sonication cycle to avoid sample heating. After sonication we added 2.6 ml of chloroform and 1.3 ml 0.5 M KCl/0.08 M H_3_PO_4_ to a final ratio of 1:2:1. Afterwards, solutions were sonicated in an ultrasonic water bath for 15 min, vortexed twice for 1 min and centrifuged for 10 min at 2.000 g to induce phase separation. The lower, hydrophobic phase was collected into a clean glass tube, dried under constant nitrogen stream, re-suspended in 500 μL of deuterated chloroform supplemented with 0.005 % BHT (Butylated hydroxytoluene) (Sigma) to prevent lipid oxidation and filled into 5 mm NMR tubes with a glass pipette.

### NMR experiments

NMR experiments were recorded on Bruker 600 MHz Avance II, and 700 MHz Avance III spectrometers equipped with triple resonance inverse NMR probes (5mm 1H/D-13C/15N TXI). All pulse sequences were from the Topspin library (Bruker, BioSpin). 1D ^1^H spectra of lipid extracts in chloroform were recorded using a pulse sequence with a 30 degree flip angle hard pulse (*zg30*), a fid resolution 0.43 Hz, 64 scans and 20 ppm SW. Processing was done using an exponential window function multiplication (EM) with line broadening (LB) of 0.1 Hz, Fourier Transform and baseline correction. 1D ^1^H spectra of live worms and leakage control spectra in aqueous M9 media were acquired using a pulse sequence with excitation sculpting for water suppression (*zgesgp*)^35^. We used a resolution of 2.72 Hz, 128 scans, and 16 ppm SW. Processing was carried out with EM (LB 5Hz) and baseline correction. For 1D ^13^C spectra, we used a pulse sequence with a 30 degree flip pulse and inverse gated decoupling (*zgig30*). Fid resolution was 1.63 Hz, NS 256 and 300 ppm SW. Processing was carried out with EM (LB 30 Hz) and baseline correction.

Full spectral range, 2D ^1^H-^13^C HSQC spectra of lipid extracts in deuterated chloroform were acquired with a phase-sensitive pulse sequence and gradient pulses (*hsqcgpph*).

We used 1K and 256 increments in ^1^H and ^13^C dimensions, respectively (resolution of 21.8 Hz and 220.1 Hz), 16 scans and 16 ppm (^1^H) and 160 ppm (^13^C) SW. ^13^C decoupling was accomplished with the GARP sequence. Processing was done using sine-bell window function multiplications, Fourier Transform and baseline correction in both dimensions. For semi-selective 2D ^1^H-^13^C HSQC’s we used the same pulse-sequence and folded signals were removed using digital quadrature detection as previously described^23^. NMR parameters for each spectral region analyzed were as follows: regions 1 and 2 (Δ-CH_2_-Δand Δ-1), 1024 and 512 increments in ^1^H and ^13^C, respectively (resolution 4.1 Hz and 2.75 Hz), 16 scans and SW of 3 ppm (^1^H, 3 ppm offset) and 4 ppm (^13^C, 27 ppm offset), experimental time 3h 15min. For region 3 (^13^C double bond area) we used 1024 and 512 increments in ^1^H and ^13^C, respectively (resolution 4.1 Hz and 2.75 Hz), 40 scans and SW of 3 ppm (^1^H, 5.4 ppm offset) and 4 ppm (^13^C, 129.5 ppm offset), experimental time was 8h 11min. Processing was done with EM (LB 5Hz) and baseline correction. Spectra of lipid extracts were collected with non-isotopically enriched samples.

Full spectral range, 2D ^1^H-^13^C HSQC spectra of live worms with uniformly, ^13^C-isotopic enrichment were acquired with a sensitivity-enhanced pulse sequence (*hsqcetgpsisp2.2*). We used 2K and 256 increments in ^1^H and ^13^C dimensions, respectively (resolution of 10.9 Hz and 206 Hz), 4 scans and 16 ppm (^1^H) and 150 ppm (^13^C) SW, experimental time was 36 min. ^13^C decoupling was achieved with the GARP sequence. Processing was done using sine-bell window function multiplications, Fourier Transform and baseline correction in both dimensions.

Real time NMR measurements to monitor UFAs signal modifications in live worms uniformly enriched with ^13^C upon temperature changes were done with the sensitivity enhanced pulse sequence (*hsqcetgpsisp2.2*) as described in the previous paragraph. We introduced the NMR tube with the live worms and started acquisitions immediately after tunning/matching and shimming. Dead-time between loading the worms into the tube and starting acquisition was taken into account for the signal build-up curves. After processing, signal intensities were extracted for selected cross-peaks and plotted against experimental time using GraphPad Prism. In all cases normalization was done with respect to the first or last spectrum of the time series. *C. elegans* metabolite specific assignments were obtained from previously published works and from the human metabolome data base^23,24,30^ (**Supplementary Table 1**). Acquisition and processing of the NMR spectra were performed using TOPSPIN 3.5 (Bruker BioSpin). 2D spectra analysis and visualization was done with Sparky^36^.

## Results and discussion

### NMR UFA analysis in C. elegans lipid extracts

We first determined overall MUFA and PUFAs composition of *C. elegans* total lipid extracts. We analyzed UFA content of the wild type strain N2 and of a *fat-3* mutant lacking the Δ6 fatty acid desaturase FAT-3^37^. This enzyme catalyzes the rate limiting step in the conversion of linoleic (LA, 18:2 n-6) and alpha-linolenic (ALA, 18:2 n-3) fatty acids (FA) into C20 PUFAs including arachidonic (20:4 n-6) and eicosapentaonoic FAs (20:5 n-3) (**Supplementary Figure 1**). Our analysis was based on a previous NMR study, which provided specific assignments for MUFAs and PUFAs in human body fluids^23^.

Comparison of 1D ^1^H spectra of N2 and *fat-3* lipid samples revealed discrete differences in spectral proton regions corresponding to unsaturated (~5.3 ppm) and saturated carbon positions (~2.38 ppm) (**Figure 1A**). As expected, NMR analysis revealed that *fat-3* worms displayed a reduction in total PUFAs content. However, the resolution accessible by ^1^H NMR did not permit the identification of individual fatty acid chains. For better resolution, we recorded 2D ^1^H-^13^C HSQC spectra on N2 and *fat-3* derived lipid extracts. While the comparative analysis of the ^1^H-^13^C HSQC of N2 and *fat-3* confirmed our previous results, we were unable to dissect changes of individual MUFA/PUFA compositions, due to poor chemical shift dispersion of long-chain FAs (**Figure 1B, C**). However, we noted that signals of olefinic carbon atoms and others in their vicinity were well-resolved, which offered the opportunity to acquire semi-selective ^1^H-^13^C HSQC spectra of defined spectral regions. Using this approach, we dramatically increased resolution and the resulting information content about these metabolites. The comparative analysis of fatty acids of N2 and *fat-3* worms clearly revealed the presence of the Δ6 desaturase substrates, 18:2 n-6 and 18:3 n-3 and an overall deficiency in C20 fatty acids, most notably virtually undetectable levels of 20:3 n-6, 20:4 n-6 and 20:5 (n-3), in full agreement with previous studies (**Figure 1D**)^37^. We detected few signals that were present in *fat-3* but absent in N2 worms (asterisk in **Figure 1D**), which suggested that *fat-3* might synthesize other non-standard PUFAs to compensate for the lack of C20 FAs, in line with recent evidence^38^. A similar analysis of *fat-4 C. elegans* mutants that lack the Δ5 desaturase enzyme and, thus, cannot synthesize 20:4 n-6 and 20:5 n-3 FAs allowed us to assign NMR signals specific for these PUFAs (**Supplementary Figure 2A, B**). Our results showed that NMR can be used to analyze MUFA and PUFA acyl chains contents in *C. elegans* extracts, providing a complementary readout to analytical methods such as gas chromatography coupled mass spectrometry (GC-MS).

**Figure 1.**
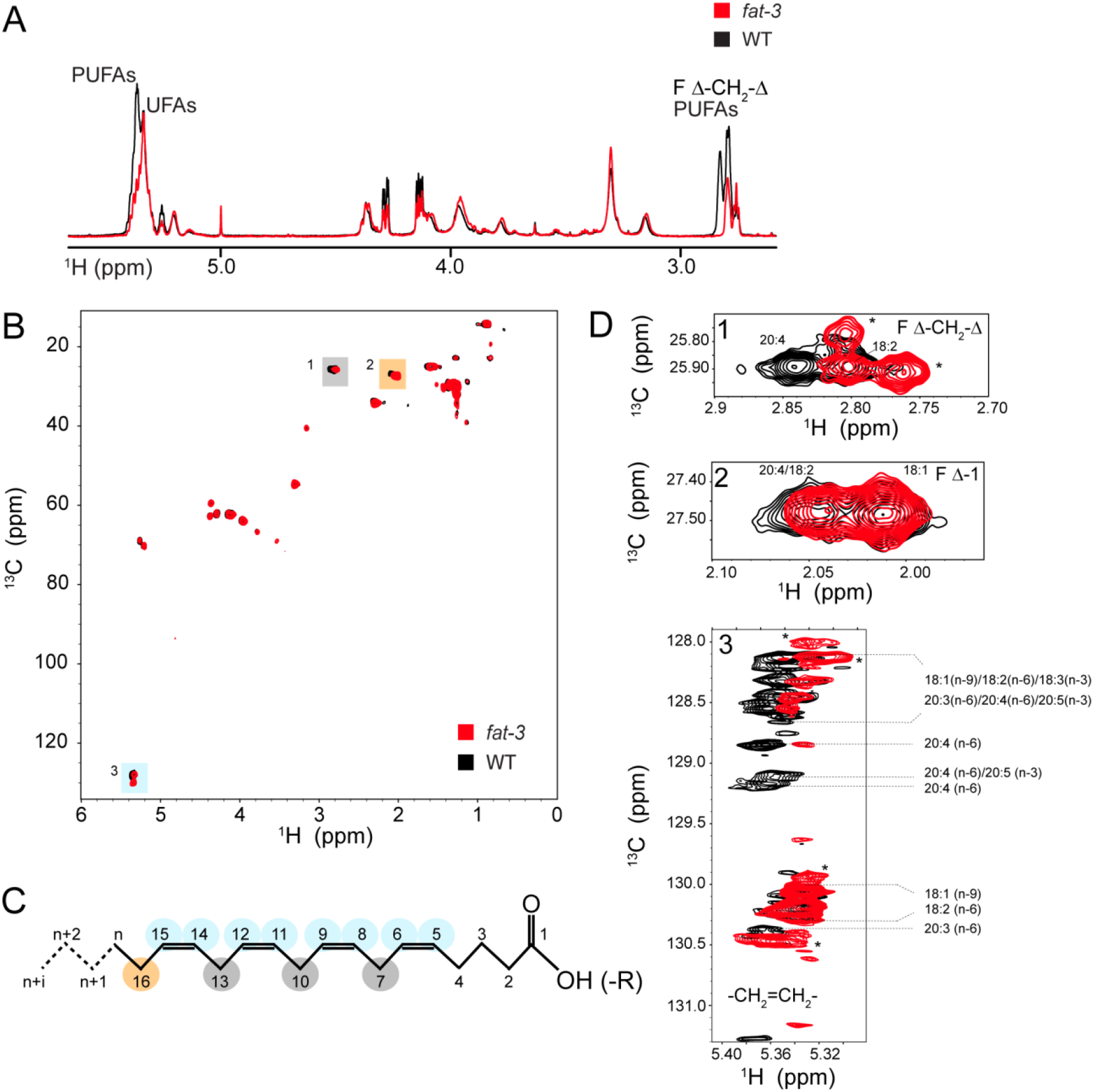
NMR analysis of *C. elegans* lipid extracts. (A) 1D ^1^H NMR spectra of lipid extracts of N2 (black) and *fat-3* (red) worms in the aliphatic region. Signals corresponding to MUFA/PUFAs are annotated. (B) ^13^C natural abundance ^1^H-^13^C HSQC spectra of N2 and *fat-3* lipid extracts. Color boxes indicate the signals of different atoms from unsaturated fatty acid side chains, as depicted in panel (C). (D) Natural abundance 2D ^1^H-^13^C semi-selective HSQC spectra for the three different regions marked in panel (B). These correspond to -CH_2_-groups between instaurations (1), -CH_2_-at position-1 from the last insaturation (2) and the double bound carbon atoms (3). Assignments from different MUFA/PUFA signals are indicated. Asterisks denote signals that appear in *fat-3* and are absent in N2 worms.

### NMR detection of UFAs in ^13^C-isotopically enriched, live C. elegans

Whereas natural abundance NMR enabled the dissection of MUFA and PUFA compositions in cell-free *C. elegans* lipid extracts, we sought to understand the dynamics of UFA metabolism in intact worms under different biological conditions. To avoid the preparation of multiple lipid extracts at individual time-points, we resorted to NMR routines on live worms. Over the past years, great progress has been made in the metabolic analysis of ^13^C-isotope enriched, intact multicellular organisms, with breakthrough studies in the water flea *Daphnia magna*^25,26^, the mold *Neurospora crassa*^39^ and more recently in *C. elegans*^30^.

We set to explore the use of NMR for the analysis of UFAs in live ^13^C-isotopically enriched *C. elegans*. Isotope labeling is stringently required to perform such experiments as field inhomogeneities due to the presence of live worms in the NMR tube results in severe ^1^H line broadening which hampers accurate signal identification and analysis using only proton NMR. In addition, while detection of natural abundance ^13^C is feasible in biological samples, the sensitivity is too low to permit the acquisition of spectra with high signal-to-noise ratios in short periods of time.

Uniform isotope labeling in *C. elegans* is a simple and inexpensive procedure that entails feeding worms with *E. coli* cultured in the presence of ^13^C-D-glucose as the sole carbon source (**Figure 2A**). Comparison of 2D ^1^H-^13^C HSQC NMR spectra of live worms uniformly enriched or at natural abundance of ^13^C showed that isotope labeling increases the overall sensitivity and allowed us to record 2D ^1^H-^13^C HSQC spectra with excellent signal-to-noise ratios in ~30 min (**Figure 2B, C**). Viability analysis during prolonged NMR acquisition times showed that most of the worms were alive and morphologically indistinguishable from control samples (**Figure 2D, E**).

**Figure 2.**
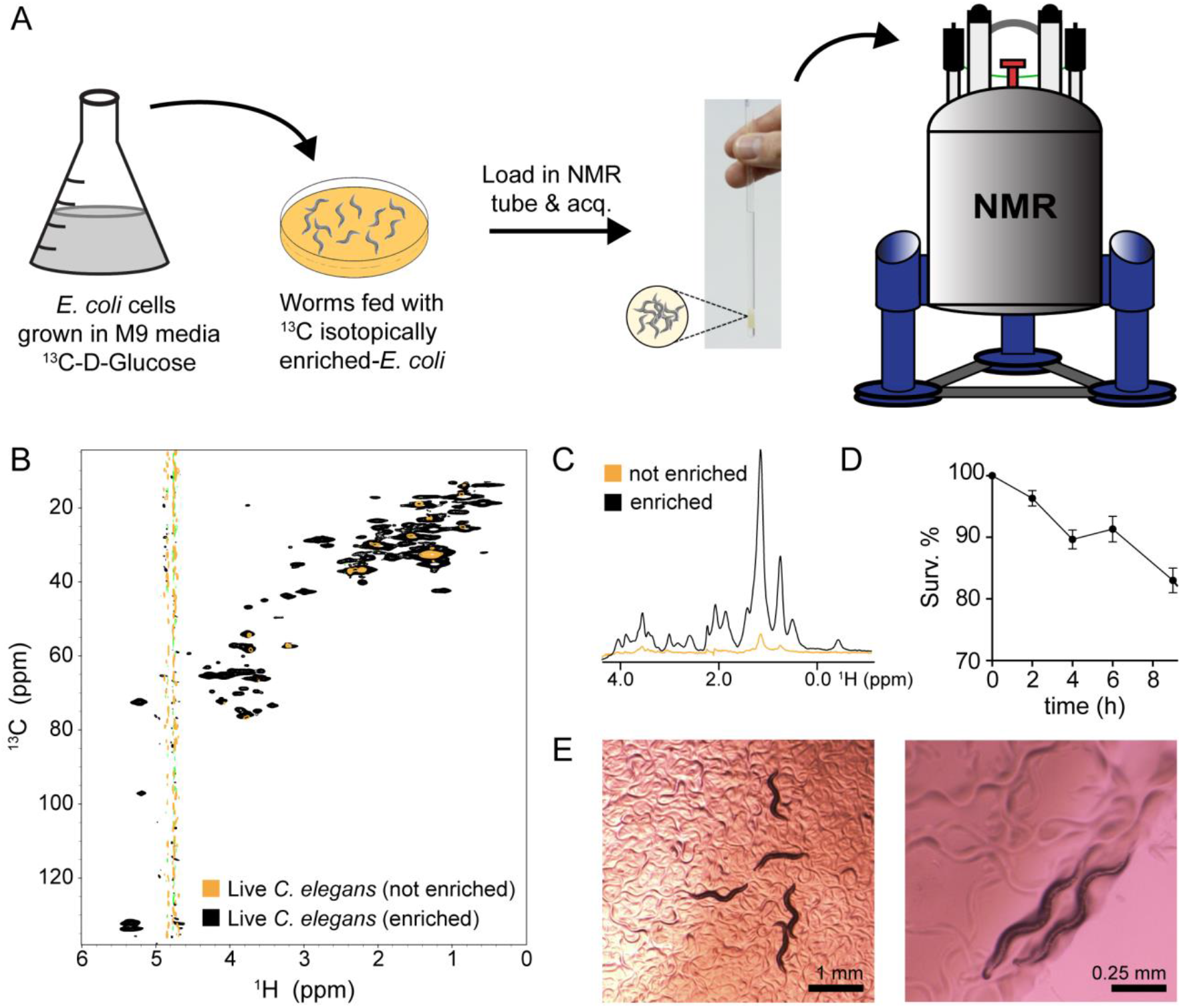
NMR analysis of live *C. elegans*. (A) Schematic representation of the protocol for recording 2D ^1^H-^13^C NMR spectra of live worms uniformly enriched with ^13^C. (B) Overlay of 2D ^1^H-^13^C HSQC spectra of non-enriched (yellow) and ^13^C uniformly enriched *C. elegans*. (C) 1D ^1^H spectra obtained from the first ser file of the spectra showed in (B). (D) Survival percentage of worms in the NMR spectrometer at different acquisition times. (E) Bright field images of worms after 4 hs at the NMR spectrometer.

To assess the reproducibility of our approach we performed replicate experiments on independent samples. As expected, increasing the number of worms in the receiver coil volume of the NMR tube resulted in proportional signal intensity increase (**Supplementary Figure 3A**). This information is useful to derive normalization factors for 2D ^1^H-^13^C HSQC analyses if one wishes to compare samples with different number of worms. Acquisition of 2D ^1^H-^13^C HSQC experiments on these replicate samples yielded almost identical spectra (**Supplementary Figure 3B**). Leakage analysis after 8 h revealed only few, low intensity peaks confirming that the bulk of NMR signals originated from live worms (**Supplementary Figure 3C, D**). The most prominently leaked metabolites were alanine, glycine, succinate, lactate and acetate. Interestingly, these metabolites were previously identified to be excretion products of the nematode ^40,41^. These observations suggested that besides its application for *in vivo* metabolic analyses, NMR can also serve as a complimentary methodology for quantitative studies of worm exo-metabolomes and respective changes under varying experimental conditions.

Next, we performed a comparative analysis with N2 and *fat-3* worms. Overlays of 1D ^13^C and 2D ^1^H-^13^C HSQC of N2 and *fat-3* worms revealed discrete changes in UFA NMR signals (**Figure 3A-C**). For *fat-3* animals, we observed a marked reduction in upfield components in the UFA double-bond spectral region (5.3 and 131 ppm in ^1^H and ^13^C dimensions, respectively). By contrast, low-field components were similar to N2 animals. As these signals correspond mostly to C18 MUFAs, whereas up-field components represent C20 PUFAs (**Figure 1**), our results suggested that *fat-3* worms were compromised in C20 PUFAs biosynthesis, in full agreement with our previous observations in lipid extracts. Apart from C20 PUFAs, we also detected changes for a small number of non-PUFA signals, indicating that *fat-3* enzymatic pathways may interconnect with other metabolic routes and/or that long-chain PUFAs also modified the biosynthesis of other compounds.

**Figure 3.**
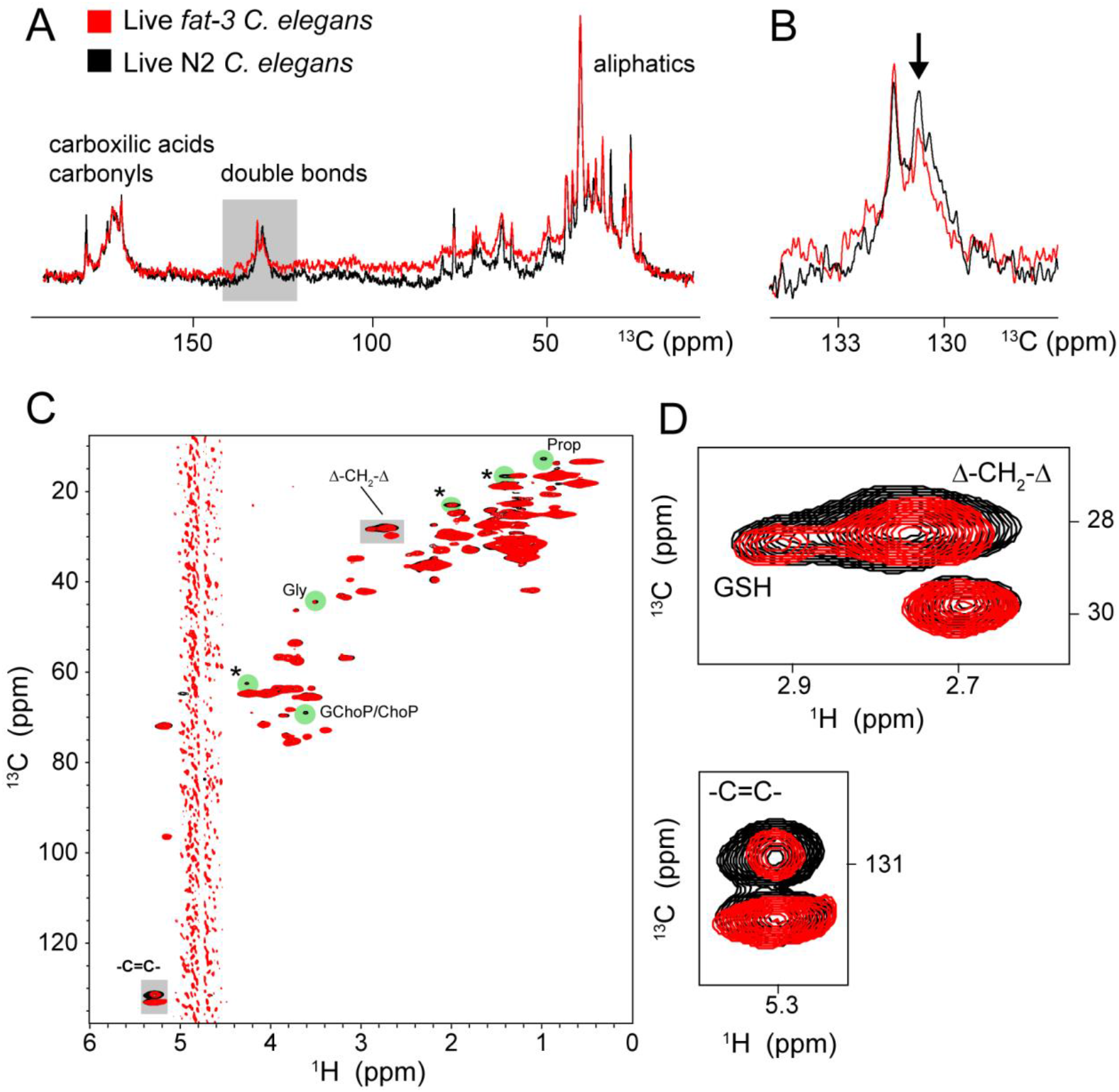
NMR characterization of UFAs in live *C. elegans*. (A) 1D ^13^C NMR spectra of ^13^C uniformly enriched N2 (black) and *fat-3* (red) *C. elegans*. The spectral region corresponding to the double-bond ^13^C atoms of UFAs is highlighted in gray and an enlarged view is depicted in (B). The arrow indicate the upfield cluster of signals in the ^13^C double-bond region corresponding to a decreased intensity of signals from long chain (20 C) PUFAs, such as arachidonic and eicosapentanoic acid. (C) 2D ^1^H-^13^C HSQC spectra of ^13^C uniformly enriched N2 (black) and *fat-3* (red) *C. elegans*. Gray boxes denote the ^13^C double-bond signals (-C=C-) and the -CH_2_-signals corresponding to carbon atoms between instaurations. (D) Enlargements of the boxed signals from panel (C) showing decreased signals of long chain PUFAs in *fat-3* nematodes. This is particularly evident for the upfield cluster of signals corresponding to the double-bonded ^13^C, in agreement with the 1D ^13^C spectra shown in (A) and (B). Green circles in panel (C) indicate major changes in metabolites other than UFAs between N2 and *fat-3*. Those that have been assigned are annotated. These correspond to: Gly (glycine), Val (valine), Prop (propionate), GChoP (glycerophosphocholine) and/or ChoP (phosphoryl choline). Asteriks indicate signals for which we don’t have unambiguous assignments.

These results further illustrated the advantages of NMR spectroscopy for studying UFAs in live *C.elegans*. The high resolution and signal-to-noise of the 2D ^1^H-^13^C NMR spectra recorded on ^13^C isotopically enriched worms suggest that the methodology may find broad applications for metabolic studies in live *C. elegans* and other multicellular organisms that can be isotopically enriched using simple growth procedures. Furthermore, the relatively short experimental times required for registering high quality 2D NMR experiments together with the little impact on the worm’s viability suggest that this approach may be used for real-time NMR UFA/PUFA analysis in live *C. elegans*.

### Real-time NMR monitoring of changes in UFA compositions in live C. elegans

Having shown that steady-state NMR measurements delineated UFA compositions in lipid extracts and intact worms, we sought to determine whether real-time NMR experiments^42^ can be used to monitor dynamic changes of UFA and PUFA profiles in live specimens. Temperature critically affects fatty acid composition in bacteria, yeast and higher organisms^43^. Temperature decreases, in particular, promote transcriptional activation of key metabolic enzymes in fatty acid biosynthesis, which results in enhanced UFA production to adjust membrane fluidity and other processes^43^.

To monitor temperature-mediated MUFA/PUFA compositions in real time in live animals, we cultivated worms at 25 °C until they reached the L4 stage, loaded them in the NMR tube and reduced the temperature to 10 °C before starting to acquire consecutive ^1^H-^13^C HSQC spectra. We monitored the evolution of UFA signals corresponding to olefinic carbons at 5.3 and 128.5 ppm (upfield signal, u) and 5.3 and 130 ppm (downfield signal, d) (**Figure 4A, B** and **Supplementary Figure 4A**). We integrated these signals and plotted them as a function of time to obtain build-up curves (**Figure 4C**). As expected, when we lowered the temperature, we observed a progressive increase in these signal intensities indicating that more MUFAs and PUFAs were present. Complimentary NMR spectra of the surrounding medium confirmed that signal intensity changes originated from UFAs inside the worms and not because of excretion into the medium (**Supplementary Figure 4B**). A comparison between u and d signals revealed that d clusters increased more rapidly than u ones. Signals within the d cluster mostly corresponded to 18:1 n-9 and 18:2 n-6, with 18:1 n-9 being the desaturation product of 18:0 catalyzed by the orthologue of the human stearoyl Δ9 desaturase Fat-7. Interestingly, expression of this enzyme is cold inducible and strongly upregulated within 3 ^1^H after transferring worms from 25 °C to 10 °C^44^. In agreement with these findings, *de novo* synthesis of 18:1 Δ9 plateaued after 3-4 ^1^H of shifting the temperature from 25 °C to 10 °C (**Figure 4A-C**). Thus, our results support a direct link between the transient induction of *fat-7*^44^ to *de novo* synthesis of 18:1Δ9 and its conversion to longer PUFAs (cluster u) within the same time interval and coinciding with fatty acid desaturation.

**Figure 4.**
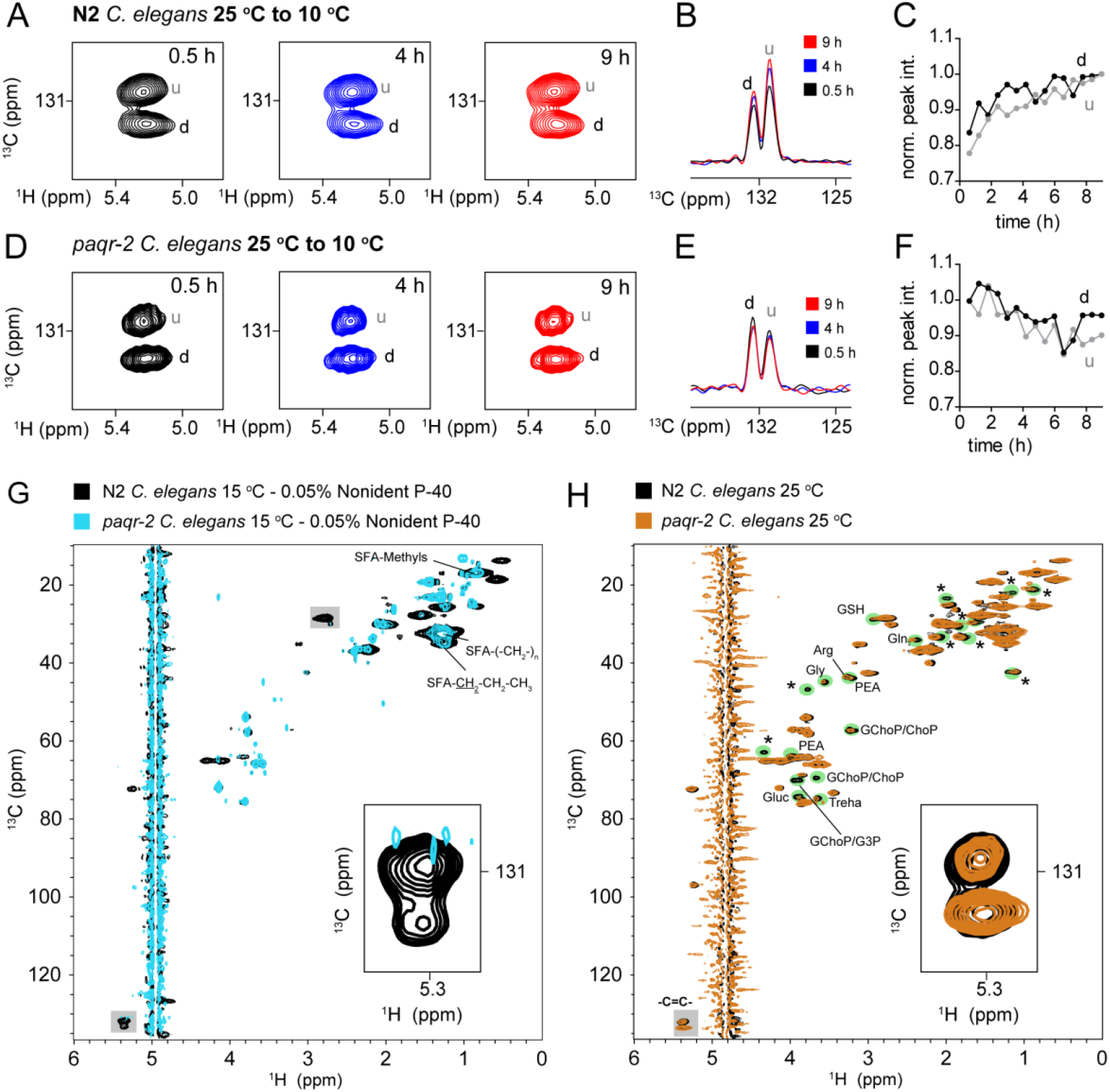
Real-time UFAs NMR analysis in live *C. elegans* during cold adaptation. Evolution of NMR signals corresponding to the double-bond ^13^C atoms of UFAs acyl chains when temperature is dropped from 25 °C to 10°C in N2 (A-C) and *paqr-2* (D-F) nematodes. 1D ^13^C NMR traces in the middle panels (B, E) show the changes in intensity for the downfield and upfield signal lobes in panels (A) and (D). Curves on the right (C, F) show the intensity of the downfield (d) and upfield (u) signal lobes as a function of time. (G) 2D ^1^H-^13^C HSQC spectra of ^13^C uniformly enriched N2 (black) and *paqr-2* (cyan) nematodes grown at 15 °C in the presence of 0.05% Nonident P-40 detergent. Gray boxes denote the UFA/PUFAs ^13^C double-bond signals (-C=C-), enlarged in the inset, and the signals corresponding to the -CH_2_-groups between instaurations. Signals from saturated fatty acids (SFA) are also indicated. (H) 2D ^1^H-^13^C HSQC spectra of ^13^C uniformly enriched N2 (black) and *paqr-2* (brown) *C. elegans*grown at 25 °C. The Gray box denote the UFA/PUFAs ^13^C double-bond signals (-C=C-) which are enlarged in the inset. Signals that show marked decreased intensities in *paqr-2* compared with N2 are indicated with green circles. These are: Gluc (β-D-glucose), Treha (threalose), ChoP (phosphoryl choline), G3P (glyceraldehyde-3P), PEA (phospho-ethanol-amine), GChoP (glycerophosphocholine), Gly (glycine), Arg (arginine), GSH (glutathione), Gln (glutamime). Asteriks indicate signals for which we don’t have unambiguous assignments.

To further study the environmental regulation of desaturation in live *C. elegans*, we repeated these experiments with the *paqr-2* mutant strain. *C. elegans* contains five proteins that belong to the PAQR family of proteins, including PAQR-1 and PAQR-2, which share high degrees of homology with mammalian adiponectin receptors AdipoR1 and AdipoR2 involved in glucose and fatty acid metabolism^13,45^. It has been shown that PAQR-2 participates in low temperature adaptation by upregulating *fat-7* expression via the nuclear receptor NH49^12,13^. Thus, PAQR2 is an essential protein for membrane adaptation to conditions promoting its rigidity, such as cold stress or diets that increase saturated fatty acids contents^46^. In *C. elegans*, the *paqr-2* mutant is compromised in its development from L1 to fertile adulthood at 15 °C^13^.

We grew *paqr-2* mutant worms at 25 °C until the L4 stage and repeated consecutive ^1^H-^13^C HSQC experiments upon shifting the temperature to 10 °C (**Figure 4D-F** and **Supplementary Fig. 4C, D**). In contrast to what we observed with N2 worms, intensity profiles of MUFAs and PUFAs signals showed that these fatty acids decreased slowly, suggesting the absence of *de novo* UFA synthesis and depletion of the existent endogenous UFA pools. These results provided further evidence in support for *de novo*MUFA/PUFA synthesis observed in N2 nematodes and suggested a direct role of PAQR-2 in regulating cold-induced UFA production to preserve membrane fluidity. To further confirm these observations, we grew *paqr-2* worms at 15 °C in the presence of 0.05 % of Nonident P-40 (NP40), a detergent that partially suppresses the cold-sensitive phenotype of the mutant, likely via increasing inherent membrane fluidity^12^. As has been shown previously, *paqr-2* worms grow at 15 °C in the presence of NP40 but do not produce progeny^10,12^. We added worms into the NMR tube and acquired 2D ^1^H-^13^C HSQC spectra, in parallel to control experiments with N2 reference animals grown under the same conditions (**Figure 4G**). We detected signals from MUFAs/PUFAs in the N2 reference sample but not in *paqr-2* worms. In contrast, signals from saturated fatty acids which serve as initial substrates for desaturating enzymes were present in both samples. These observations, confirm the pivotal role of PAQR-2 in the activation of Fat-7 followed by UFA biosynthesis at low temperatures.

Next, we analyzed NMR signals from molecules other than MUFA/PUFAs in N2 and *paqr-2* nematodes shifted from 25 °C to 10 °C (**Supplementary Figure 4 and Supplementary Figure 5A**). In both samples, we detected rapid depletion of sugar signals corresponding to β-D-glucose and threalose, suggesting that these became primary carbon sources under conditions that favored fermentation over aerobic respiration. In parallel, we observed a substantial increase in acetate, lactate, succinate, alanine and glycine NMR signal intensities, which resulted from excretion, as discussed previously. Interestingly, signals from saturated fatty acids displayed differential behaviors between N2 and *paqr-2* specimens upon cold adaptation (**Supplementary Figure 5B-D**). As substrates of Δ9-desaturases, saturated fatty acid acyl-chain signal intensities increased in N2 samples during the first 3-4 h of cold adaptation, until they reached steady-state levels. This time-frame agrees well with the highest transcriptional induction of *fat-7*^44^ and the progressive *de novo* biosynthesis of UFAs (**Figure 4 A-C**). By contrasts, we only detected a slight, initial increase of saturated FA signal intensities in *paqr-2* worms, which eventually decreased and plateaued after ~5 h. This is consistent with the mutant worm being unable to desaturate *de novo* synthesized FAs during cold adaptation and them being metabolized.

Finally, we compared 2D ^1^H-^13^C HSQC spectra of N2 and *paqr-2* worms at 25 °C to identify additional, PAQR2-dependent metabolic changes (**Figure 4H**). We detected discrete alteration of metabolite signals other than MUFAs/PUFAs, which insinuated that *paqr-2* mutations may exert additional effects on other pathways. Importantly, we identified attenuations of phosphoethanolamine (pEA) and phosphocholine (pCho)/glycerophosphocholine (GpCho) NMR signal intensities in *paqr-2* strains. These metabolic intermediates are key constituents of the methylation-dependent *de novo* phosphatidylcholine (PC) biogenesis pathway in *C. elegans*^47^. Accordingly, genetic mutations that limit PC production have been shown to increase SBP-1-dependent expression of the *fat-7* desaturase^47^. Together, these observations suggest that impairment of the methylation pathway in *paqr-2* may explain the partial rescue bestowed by *fat-7*^12^ expression at permissive temperatures, thus enabling growth at temperatures where membrane fluidity is not fully compromised. In agreement with this hypothesis, *paqr-2* phenotypes can be rescued by blocking PC synthesis, which leads to a concomitant increase in *fat-7* expression^12^. We rationalize that a link between PAQR-2 and methylation-dependent PC biogenesis exists in *C. elegans*, however additional genetic, biochemical and metabolic experiments are needed to confirm this notion.

## Conclusions and perspectives

In this work, we used high-resolution heteronuclear NMR methods to analyze the composition of UFAs in lipid extracts and in ^13^C isotopically enriched live *C. elegans*.NMR is non-destructive, provides high-resolution structural information of biomolecules and is quantitative, thus permitting the identification and quantification of relative concentrations of metabolites in a time-resolved manner. We monitored UFA biosynthesis in real time in live worms during cold adaptation, a well-known phenomenon that results in accumulation of unsaturated fatty acids^44,48^. Our results are in full agreement with the time frame of fatty acid desaturase activation and cellular production of MUFAs and PUFAs during cold adaptation^11,12,44^. Moreover, we confirmed the pivotal role of *paqr-2* in maintaining membrane fluidity at low temperatures. In comparison to other analytical techniques for studying fatty acids in biological samples, such as HPLC-GC MS, our NMR approach has certain limitations. On one hand, the intrinsic low sensitivity of NMR spectroscopy restricts analyses to most abundant species. On the other hand, sparsity of data and NMR signal overlap compromise exhaustive detection and identification of all lipid components in a sample, and whether lipid signals originate from free fatty acids, lipid-rich particles or membranes. Given that membrane-embedded lipids exhibit severely restricted rotational motions, we do not expect them to contribute to solution-state NMR spectra. Therefore, it is more likely that detected fatty signals correspond to molecules that tumble freely in their respective environments. Lacking specialized adipocytes, *C. elegans* store most fatty acids in non-membrane-bound lipid droplets, which are highly abundant, heterogeneous organelles in worms^15^. It seems reasonable to assume that *in vivo* NMR signals of FAs originate from these particles.

Despite these limitations, our approach has equally enticing advantages because it enables a position-dependent assessment of fatty acid unsaturation levels and to determine real-time dynamics of fatty acid interconversions in response to altered environmental conditions such as cold adaptation. These features provide unparalleled insights into the activities and regulation of cellular, fatty acid-modifying enzymes. In our case, we used discrete temperature modulations to investigate effects on *in vivo* lipid metabolism. Our strategy may find additional applications in studying dietary regimes, pharmacological treatments and/or the use of genetic mutants that influence lipid homeostasis. Many of the open questions in lipid-metabolism research are directly linked to human diseases and analytical methods to address these questions are in great demand.

It should be stressed that this strategy is not solely for lipid analysis but may have broad applications to other metabolites. If assignments are available for the metabolites of interests their relative concentrations and variations upon different treatments/genetic backgrounds can be easily assessed. This is nicely illustrated in a recent paper where the metabolism of sugars, amino acids and other small compounds was profiled in ^13^C isotopically enriched N2 *C. elegans* and AAK/AMP mutants with differential carbon usage^30^. Furthermore, the introduction of NMR bioreactors and flow-cells inside the NMR tube considerably expands the scope of these applications as it allows for a precise control of energy sources and different *in situ* treatments on live multicellular organisms that may perturb their metabolism^28^. With the advent of comprehensive metabolite NMR databases such as the human metabolome database^24^, new developments in NMR spectrometers aided to increase spectral resolution and signal-to-noise and selective labeling schemes designed to feed specific metabolic routes we foresee that the high resolution NMR analysis of live organisms will provide a much deeper understanding of complex metabolic pathways.

## Acknowledgements

This work was funded by CONICET, ANPCyT (PICT-2018-02572) and the Richard Lounsbery Foundation. We acknowledge Philipp Selenko for reading and commenting on the manuscript, Andrea Coscia and Alejandro Gago for maintenance of the NMR infrastructure and Cecilia Vranych for assistance with *C. elegans* growth and maintenance.

