## Supporting Information for "Unsaturated fatty acids profiling in live *C. elegans* using real-time NMR spectroscopy"

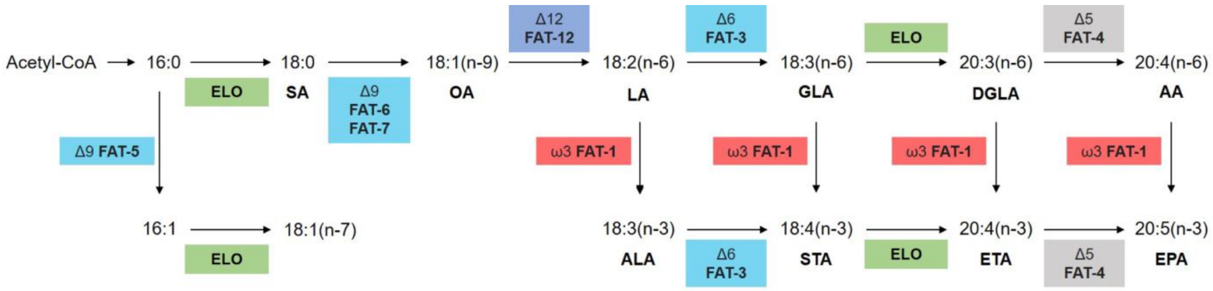


**Supplementary Figure 1.** Biosynthesis of omega-6 and omega-3 fatty acids in *C. elegans*. Unlike most other animals, *C. elegans* possesses Δ12 desaturase and Δ6 desaturase enzymes. Enzyme names and activities are enclosed in squares. Abbreviations: ELO, elongase; SA, stearic acid; OA, oleic acid; LA, linoleic acid ALA, alpha linoleic acid; GLA, gamma linoleic acid; STA, stearidonic acid; DGLA, dihommo gamma linoleic acid; ETA, eicosatetraenoic acid; AA, arachidonic acid; EPA, eicosapentaenoic acid.

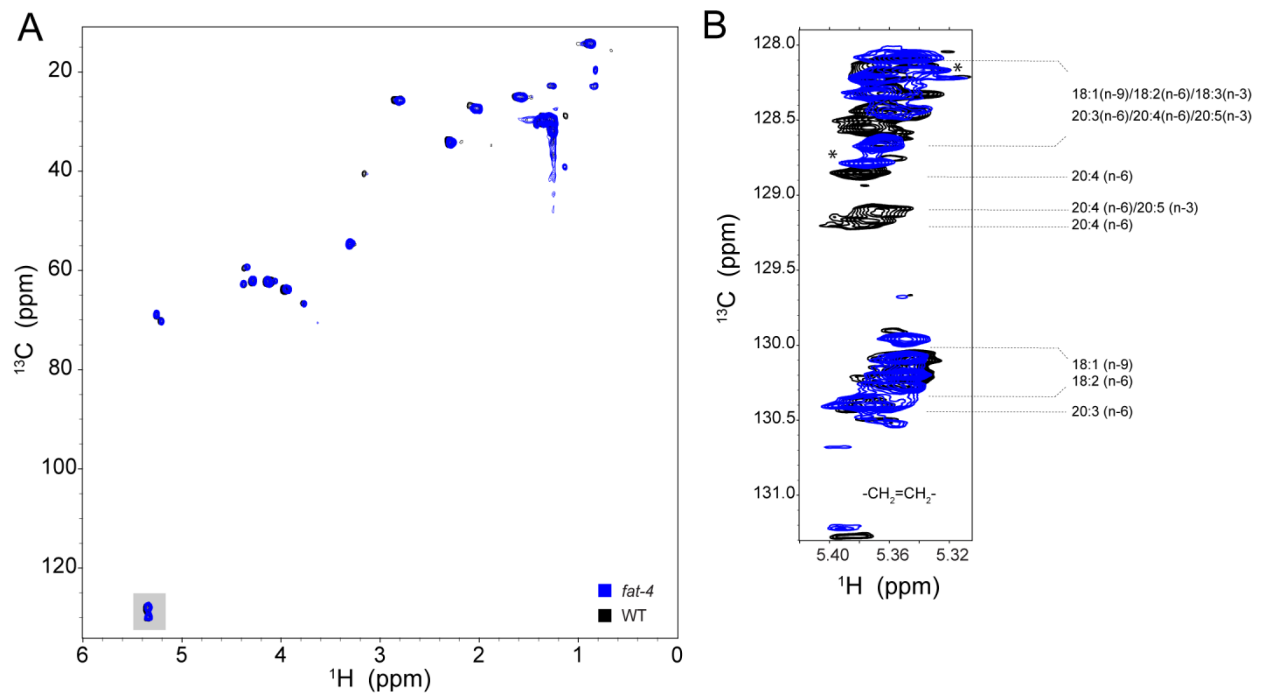


**Supplementary Figure 2.** NMR analysis of *C. elegans* lipid extracts. (A) Natural abundance 2D $^{1}H$-$^{13}C$ HSQC spectra of N2 and *fat-4* lipid extracts. (B) Natural abundance 2D $^{1}H$-$^{13}C$ semi-selective HSQC spectra of the double-bond region marked with a gray box in panel (A).

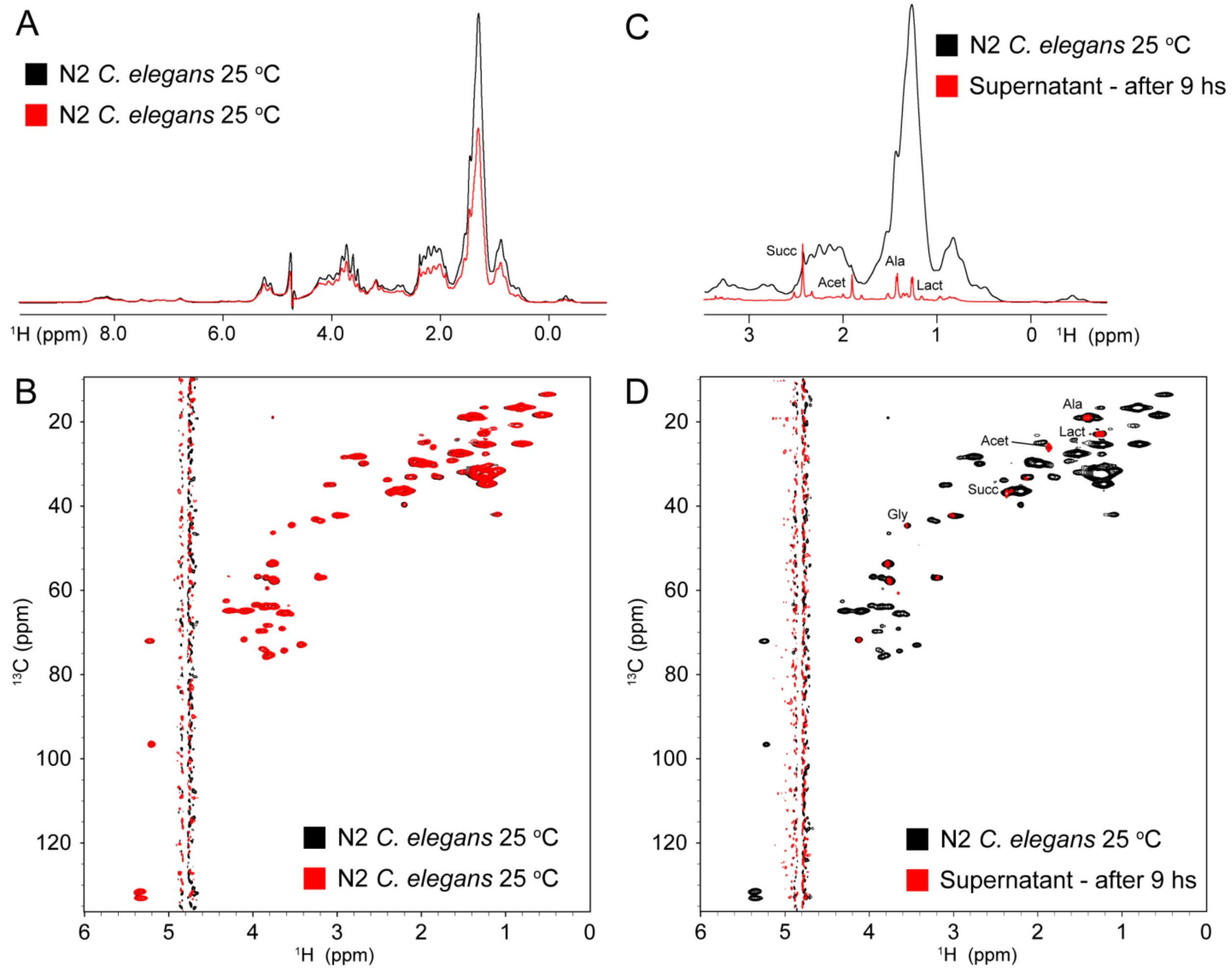


**Supplementary Figure 3.** Reproducibility of live worms NMR experiments. (A) 1D $^{1}$H NMR spectra of two independent N2 worm samples. The differences in intensity correlates with the different number of worms loaded in the NMR tube for each determination. These were ~40000 and ~60000 for the red and black spectra, respectively. (B) 2D $^{1}$H-$^{13}$C HSQC spectra of the same two samples. For comparative purposes, we multiplied the black spectra by a factor of 0.6 obtained from the comparison of the 1D $^{1}$H analysis. (C and D) Leakage controls. 1D $^{1}$H (C) and 2D $^{1}$H-$^{13}$C HSQC (D) spectra of live worms (black) and the surrounding buffer separated after 9 hs of NMR acquisitions. Leaked or excreted metabolites for which we have unambiguous assignments are indicated. Gly (glycine), Succ (succinate), Acet (acetate), Lact (lactate), Ala (alanine).

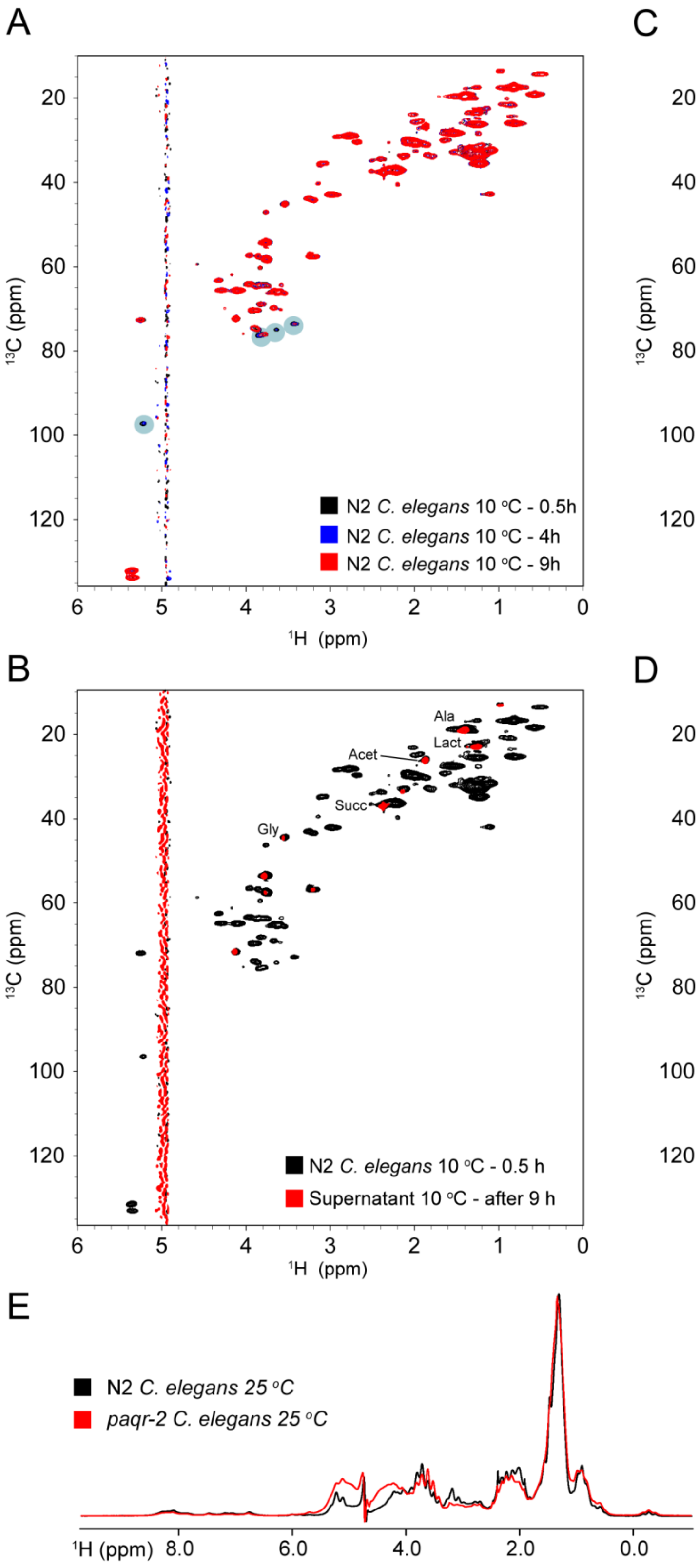
A
B
C
D
E
$^{13}$C (ppm)
$^{1}$H (ppm)
N2 C. elegans 10 °C - 0.5h
N2 C. elegans 10 °C - 4h
N2 C. elegans 10 °C - 9h
paqr-2 C. elegans 10 °C - 0.5h
paqr-2 C. elegans 10 °C - 4h
paqr-2 C. elegans 10 °C - 9h
Ala
Lact
Acet
Succ
Gly
N2 C. elegans 10 °C - 0.5 h
Supernatant 10 °C - after 9 h
paqr-2 C. elegans 10 °C - 0.5 h
Supernatant 10 °C - after 9 h
N2 C. elegans 25 °C
paqr-2 C. elegans 25 °C
$^{1}$H (ppm) 8.0 6.0 4.0 2.0 0.0

**Supplementary Figure 4.** Real-time NMR analysis of $^{13}C$-isotopically enriched N2 and *paqr-2* worms. 2D $^{1}H$-$^{13}C$ HSQC spectra of N2 (A) and *paqr-2* (C) worms at different time points after the temperature was dropped from 25 $^{o}C$ to 10 $^{o}C$. Signals marked in cyan correspond to rapidly metabolized sugars, including glucose and threalose. (B and D) Leakage controls. 2D $^{1}H$-$^{13}C$ HSQC spectra of live N2 (B) and *paqr-2* (D) worms (black) and the surrounding buffer separated after 9 hs of NMR acquisitions at 10$^{o}C$ (red). (E) Comparative 1D $^{1}H$ NMR spectra of N2 (black) and *paqr-2* (red) $^{13}C$ isotopically enriched, live worms.

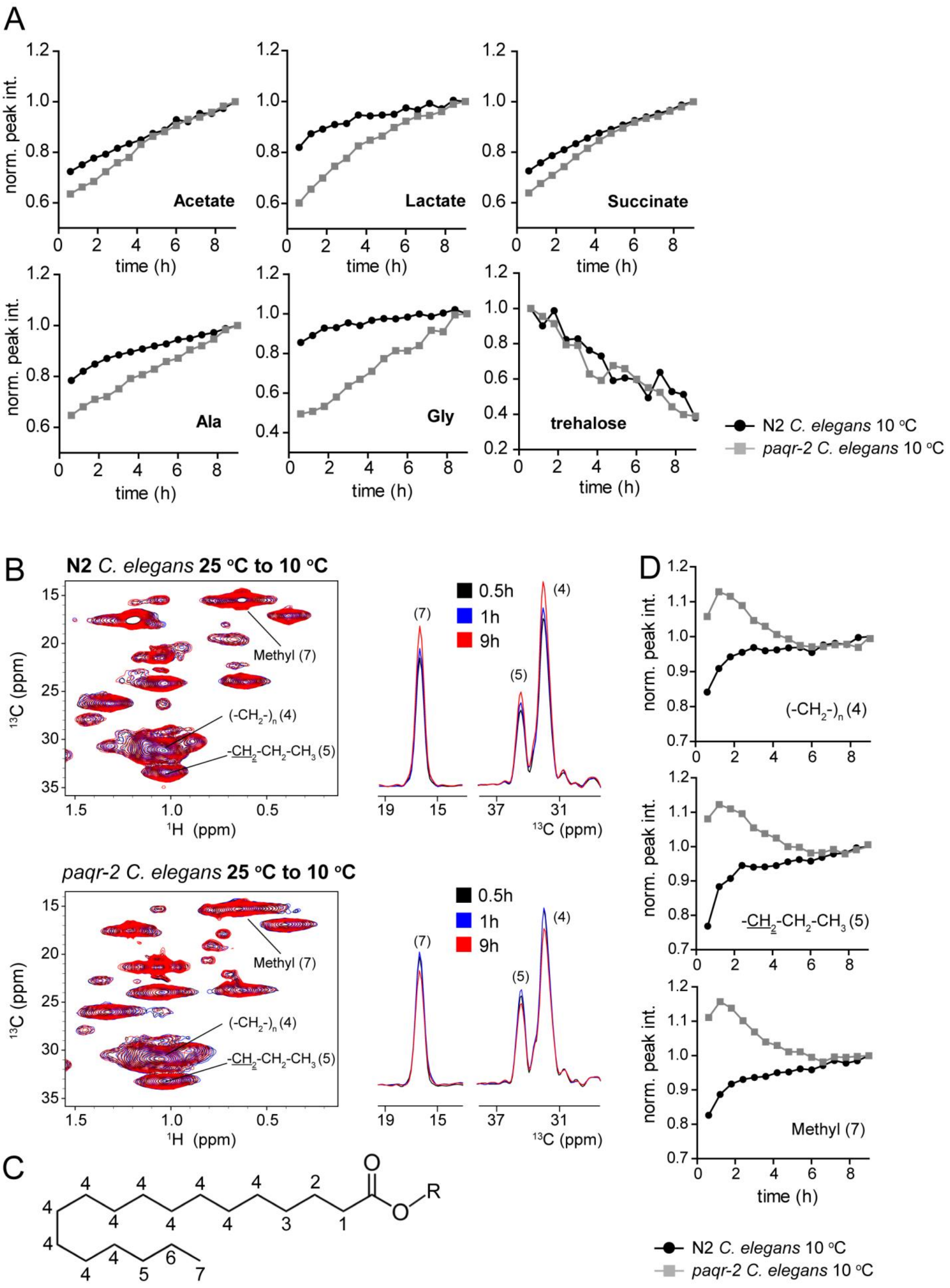

A
Acetate
Lactate
Succinate
Ala
Gly
trehalose
norm. peak int.
time (h)
N2 C. elegans 10 °C
paqr-2 C. elegans 10 °C
B
N2 C. elegans 25 °C to 10 °C
paqr-2 C. elegans 25 °C to 10 °C
Methyl (7)
(-CH2-)n (4)
-CH2-CH2-CH3 (5)
13C (ppm)
1H (ppm)
0.5h
1h
9h
C
O
R
D
(-CH2-)n (4)
-CH2-CH2-CH3 (5)
Methyl (7)
norm. peak int.
time (h)
N2 C. elegans 10 °C
paqr-2 C. elegans 10 °C

**Supplementary Figure 5.** Real-time NMR analysis of different metabolites in live *C. elegans*. (A) Evolution of NMR metabolite signals when temperature is dropped from 25 $^{o}$C to 10$^{o}$C in N2 (black) and *paqr-2* (gray) nematodes enriched with $^{13}$C. Signal intensities were extracted from sequential $^{1}$H-$^{13}$C HSQC spectra after decreasing the temperature. (B) 2D $^{1}$H-$^{13}$C HSQC spectra (left) and 1D $^{13}$C traces (right) at frequencies corresponding to different atoms from saturated fatty acid chains in N2 (top) and *paqr-2* (bottom) nematodes when temperature is dropped from 25 $^{o}$C to 10 $^{o}$C . (C) Schematic representation of a saturated fatty acid chain. Carbon atoms with distinct chemical shift values are identified. (D) Signal intensity as a function of experimental time for the signals indicated in (B) when the temperature is decreased.

**Supplementary table 1.** Chemical shifts values of NMR signals from $^{1}H$-$^{13}C$ HSQC spectrum of live worms.[a]

| Compounds | Chemical group | $^{13}C$ | $^{1}H$ |
|---|---|---|---|
| Acetate | Methyl | 26.12 | 1.9 |
| Alanine | CβHβ | 19.06 | 1.449 |
| Arginine | CδHδ | 43.04 | 3.243 |
| Glycerophosphocholine | Methyl | 56.739 | 3.19 |
| Glycerophosphocholine | $CH_2$(2) | 68.72 | 3.636 |
| Glycerophosphocholine | $CH_2OH$ | 69.472 | 3.881 |
| Phosphorylcholine | Methyl | 56.741 | 3.185 |
| Phosphorylcholine | $CH_2$(2) | 68.77 | 3.662 |
| Glutathione | Cysteinyl CβHβ | 28.35 | 2.927 |
| Glutamine | CγHγ | 33.733 | 2.426 |
| Glycine | CαHα | 44.44 | 3.54 |
| Lactate | Methyl | 22.915 | 1.309 |
| Propionate | Methyl | 12.94 | 1.037 |
| Succinate | $CH_2$ | 36.84 | 2.385 |
| α-D-Glucose | $C_1H_1$ | 94.314 | 5.211 |
| Saturated fatty acids | $\underline{CH_2}$-$CH_2$-$CH_3$ (5) | 34.762 | 1.264 |
| Saturated fatty acids | Methyl (7) | 16.64 | 0.867 |
| Saturated fatty acids | $(CH_2)_n$- (4) | 32.232 | 1.298 |
| β-D-Gluc | $C_1H_1$ | 98.74 | 4.629 |
| β-D-Gluc | $C_2H_2$ | 77.064 | 3.213 |
| β-D-Gluc | $C_3H_3$ | 78.694 | 3.448 |
| Trehalose | $C_1H_1$ | 96.031 | 5.163 |
| Trehalose | $C_2H_2$ | 73.9 | 3.63 |
| Glyceraldheyde 3-P | $CH_2OH$ | 69.34 | 3.866 |
| Phosphoethanolamine | $CH_2$-N- | 43.48 | 3.19 |
| Phosphoethanolamine | $CH_2$-O-P | 63.138 | 3.944 |
| Unsaturated fatty acids | Δ-1 | 29.855 | 2.075 |
| Unsaturated fatty acids | Δ-$CH_2$-Δ | 28.235 | 2.8 |
| Unsaturated fatty acids | -C=C- (upfield set) | 130.65 | 5.31 |
| Unsaturated fatty acids | -C=C- (downfield set) | 132.241 | 5.289 |

[a]Chemical shifts values are reported at 20°C. Assignments were obtained from Nguyen et al, Anal. Chem. 92, 7382-7387, (2020); Willker and Leibfritz Mag. Res. Chem. 36, S79-S84, (1998) and the Human Metabolome database (https://hmdb.ca).
